# Modifying Azithromycin to Mitigate Arrested Autophagy

**DOI:** 10.1101/2024.04.25.591217

**Authors:** Ryan D Quarrington, Shiye Chen, Sylvia A Sapula, Shawn W H Liu, Susan E Lester, Matthew M Miller, Vesna Munic Kos, Benjamin T Kopp, Hubertus PA Jersmann, Anton Blencowe, Eugene Roscioli

## Abstract

**Aims/hypothesis:** Autophagy plays a critical role in the survival and microbial clearance function of tissues that encounter the environment, such as airway epithelial cells (**AEC**s). Contrary to the known roles of azithromycin (**AZM**) in promoting microbial clearance, our evidence shows that AZM is a potent inhibitor of autophagy ‒ an effect consistent with bacterial residency. Here we investigate the structure-activity relationship of AZM *vs* other macrolides and AZM-3’-*N*-oxide (**AZM-[O]**), to mitigate the off-target arrest of autophagy.

**Method:** AZM-[O] was synthesised in-house via selective oxidation of the desosamine amine of AZM. Human peripheral blood mononuclear cells (**PBMC**) were used to assess autophagy *ex vivo* in an organotypic manner via transmission electron microscopy and Western blot analysis. For *in vitro* studies, the 16HBE14o-AEC line and Western blot was used to assess macrolide *vs* autophagy structure-activity relationships, and autophagic flux by quantifying the protein abundance of LC3B-II *vs* Sequestosome-1. Subsequent assessments of antimicrobial activity were conducted using the micro-broth dilution method. Immunomodulatory outcomes were assessed by quantifying the secretion of IL-6 in a lipopolysaccharide PMA-stimulated THP-1 macrophage model.

**Results:** AZM significantly inhibited autophagic flux in both the *ex vivo* PBMC and *in vitro* 16HBE14o-AEC models, evidenced by the accumulation of autophagosome-related vacuoles and LC3B-II and Sequestosome-1 protein, compared to its precursors and other macrolides including roxithromycin and clarithromycin. Notably, oxidation of AZM to produce AZM-[O] significantly alleviated this inhibitory effect on autophagy, but without completely preserving its antimicrobial and immunomodulatory functions.

**Conclusion/Directions:** We show for the first time that AZM inhibits PBMC autophagy in human blood *ex vivo* and there is a high probability this phenomenon occurs in the clinical setting. Importantly, chemical modification of AZM to generate AZM-[O] substantially alleviated this effect, consistent with altered ionisation properties. We are now assessing clinically derived blood samples in participants treated with AZM, defining the AZM–mammalian protein interactome and developing further AZM derivatives that preserve immunomodulatory activity while minimising disruption of autophagy.

## Introduction

The consequences of intracellular bacteria surviving for prolonged periods in individuals who are routinely exposed to our most significant therapeutics captures some of the world’s most immediate health priorities: the expansion of bacterial species that simultaneously adapt to exploit the intracellular niche and our frontline antibiotics. Accordingly, a great proportion of the world’s population live for years with respiratory conditions that require antibiotic therapy (e.g., cystic fibrosis, chronic obstructive pulmonary disease (**COPD**) and bronchiectasis). For example, despite the broad window to treat COPD, this condition remains in the top three leading causes of mortality (1, 2), and recurrent microbial infection that can reside in long-lived airway epithelial cells (**AECs**) continue to be the primary driver of disease (3-8). Moreover, macroautophagy (hereafter referred to as autophagy), the principal mechanism responsible for clearing intracellular microbes (9), is frequently dysregulated in pulmonary disease (10, 11). Hence, the pulmonary surface in individuals with chronic respiratory disease represent a vast reservoir for bacterial adaptation (12). The convergence of these phenomena coupled with the reliance on antibiotics (to prevent a sudden increase in global mortality (13)), may be the central reason that despite multiple decades of pharmacological innovation (14), current therapeutic options are palliative and airway infections remain central to disease progression. It is critical to safeguard the efficacy of our frontline antibiotics in the absence of therapeutic options that target the causal mechanisms of pulmonary disease.

As for any medical intervention directed against a microbial pathogen, antibiotics can also elicit undesirable off-target effects in mammalian cells. While it is reasonable to justify this when the off-target effects are well tolerated, a greater concern is enhancing biochemical disease mechanisms, that are beyond the sensitivity of diagnostic/clinical inquiry. A prime example of this would be blocking the host-cell’s innate bacterial clearance mechanisms. We and others have found that azithromycin (**AZM**) likely blocks autophagy (4, 15, 16). Normally, in the absence of disease, bacteria are encapsulated within autophagosomes and are rapidly eliminated upon lysosomal fusion by digestive enzymes and acidic pH (9). Indeed, the consequences of perturbing the many other autophagy-mediated functions align with remarkable fidelity to the hallmark phenotypes ascribed to pulmonary disease, such as accelerated senescence (17), nutrient deprivation (18), uncontrolled autoimmune-like inflammation (19), dysfunctional cilia (20), and mitochondrial dysfunction (21). The interplay between autophagic dysregulation and respiratory disease is expansive and there have been several attempts to effectively describe their relationship (11, 22-24). One reason this off-target effect by AZM has been left unchecked is because clinical assessments of infection are confined to symptom relief, and methods in diagnostic pathology are not geared to detect intracellular bacteria. Furthermore, to our knowledge, autophagy is completely disconnected from clinical and diagnostic inquiry. Setting aside the fact that autophagy is an influential survival mechanism, this points to a scenario whereby airway cells have a heightened vulnerability to persistent intracellular bacteria due to restricted xenophagy. Hence, it is important to determine how this occurs and whether AZM can be modified to maintain its efficacy and allow the cell’s autophagic functions to operate normally.

AZM is a third-generation semisynthetic macrolide and is distinct from its parent molecule erythromycin A (**ERM**), due to expansion of the lactone ring through insertion of a methylated nitrogen atom ( (25); and see **Fig. 2A** for macrolide structures). The resulting enhanced pharmacokinetics, spectrum of activity and tolerance of AZM (*vs* ERM) has made it one of the world’s most frequently prescribed antibiotics, which also comports with significant stewardship issues (26, 27). In part owing to its lipophilicity, AZM concentrates at the vast surface area afforded by the ciliated airway epithelium and within the airway surface liquid (25, 28). Coupled with this, AZM’s physiochemical properties exhibit a remarkable capacity to passage pH and electrochemical gradients established by cellular and intracellular membranes (29). As lysosomes partition the lowest pH environment in mammalian cells, this makes AZM highly lysosomotropic. We and others have shown AZM accumulates in lysosomes via non-ionic diffusion at levels orders of magnitude higher than the cytosol or extracellular matrix (30-32). Aside from being sequestered from microbes in the cytosol, AZM increase lysosomal pH via ion trapping due to the protonation of two of its basic functional groups. This prohibits the normal function of lysosomal proteases, blocks autophagosome-lysosome fusion and inhibits autophagic flux with similar efficacy as bafilomycin (a widely applied inhibitor of autophagy) (4, 16). Indeed, a recent systematic review comparing intracellular antibiotic activity reported that AZM ranked among the lowest-performing agents for clearance of intracellular *Staphylococcus aureus* in experimental models of osteomyelitis (33). What is more, the gradual leakage of AZM from lysosomes exposes bacterial communities to sub-lethal amounts of the antibiotic, enhancing their intracellular persistence (5). The importance of addressing this scenario for AZM, autophagy and chronic respiratory disease, is further underscored by current concerns surrounding the paucity of new antibiotics entering the drug development pipeline (34, 35).

Given these significant challenges, we assessed the structure–activity relationship (**SAR**) of AZM in relation to autophagic flux. These outcomes were compared with other clinically relevant macrolide antibiotics and azithromycin-3′-N-oxide (**AZM-[O]**), a partially oxidised AZM derivative designed to counter lysosomal protonation. Autophagy, inflammatory, and antimicrobial outcomes were assessed in models relevant to the clinical administration of AZM and the human airway, a major site of therapeutic exposure in chronic respiratory disease.

## Methods

### Synthesis of Azithromycin-3’-*N*-Oxide

AZM-[O] was prepared by adapting two previously described methods (36, 37). AZM (100 mg, 134 µmol) was dissolved in methanol (1 mL) in a single necked round bottomed flask (5 mL) and cooled to 0 °C with an ice bath. 30% w/v H_2_O_2_ (200 µL, 1.96 mmol) was added dropwise over 30 min and the reaction was stirred at 0 °C for 90 min. The mixture was added to ice water (20 mL) and dichloromethane (10 mL) and stirred at 500 rpm for 1 min. A saturated sodium thiosulfate solution (15 mL) was added, and the mixture was transferred to a separating funnel. The organic phase was removed and the aqueous extracted with dichloromethane (2 × 10 mL). The combined organic extracts were dried with magnesium sulfate and filtered under vacuum. The filtrate was concentrated *in vacuo* and dried (0.01 mbar, 22 °C, 16 h) to afford the title compound as a white solid, 86 mg (84%). ESI-MS [M+H]^+^ calculated = 765.51 m/z; measured = 765.62 m/z. Reagents were purchased from ChemSupply (SA, Australia), except for AZM, which was purchased from Sigma-Aldrich.

### Biological Models

*Ex vivo* analysis of autophagy in human peripheral blood mononuclear cells (**PBMCs**) was performed as previously described (38), with the exception that blood (per participant) was also exposed to 50 µg/mL pharmacological grade AZM (AFT Pharmaceuticals Pty Ltd) for increasing time periods, in addition to the no-treatment (**NT**) and chloroquine diphosphate (**CQ**; Sigma Aldrich, C6628; 150 µM) conditions. Peripheral blood was obtained by the clinical staff of South Australia Pathology. Participants were free of current illness or chronic disease (n = 4, age 31.5 +/-11.3). Ethics approval for the use of human tissues was granted by the Central Adelaide Local Health Network Human Research Ethics Committee (study reference number: 12978). Methods involving human samples were conducted in accordance with the Declaration of Helsinki, and with the understanding and the written consent of each participant.

**16HBE14o-** (<u>h</u>uman <u>b</u>ronchial airway <u>e</u>pithelial cells) were purchased from EMD Millipore Corporation (Temecula, CA, USA) and propagated as per the manufacturers’ instructions. Culture flasks were coated with human fibronectin (10 µg/mL), bovine serum albumin (Fraction V Stock, 100 µg/mL), and PureCol (collagen, 30 µg/mL) suspended in α-MEM (each Sigma-Aldrich, MI, USA). 16HBE14o-expansion media was α-MEM (Sigma) supplemented with 10% fetal calf serum, 1× penicillin/streptomycin and 2 mM L-glutamine (GlutaMAX; each Thermo Fisher Scientific, VIC, Australia). Cells were grown in standard mammalian cell culture conditions at 37°C and 5% CO_2_, with humidity. Endpoint culture formats were 12-well plates (Corning, NY, USA), seeded with 2 × 10^5^ cells/well in 2 mL of growth media.

The **THP-1** peripheral blood monocyte cell line was purchased from the American Type Culture Collection (Manassas, VA, USA). Growth and differentiation conditions were as previously described (39), with modifications to support conditions that approximate the proteomic profile observed in human macrophages (40). Standard mammalian cell culture conditions were used (37°C and 5% CO2, with humidity), and cells maintained in RPMI 1640 supplemented with 50 µM ß-mercaptoethanol (both Sigma-Aldrich), 10% fetal calf serum, 1× penicillin/streptomycin and 2 mM L-glutamine (as above). Macrophage differentiation was performed for 2 × 10^5^ cells/well (12-well plate format), by supplementing the growth media with 50 ng/mL phorbol-12-myristate-13-acetate (PMA, Sigma-Aldrich) for 16 h, followed by a 48 h rest period.

The *in vitro* models were exposed to AZM-[O], solithromycin (**SLM**; Cayman Chemicals, MI, USA), AZM, erythromycin A (**ERM**), telithromycin (**TEL**), roxithromycin (**ROX**), clarithromycin (**CLM**; each Sigma-Aldrich), erythromycin-9-oxime (**ERM-oxime**) and erythromycin-9,11-imino ether (**ERM-ether**; Toronto Research Chemicals, ON, Canada), dissolved in molecular biology grade dimethyl sulfoxide (DMSO, Sigma-Aldrich) to less than 0.1% v/v in media. Chemical structures of the macrolides were generated using ChemDraw v23.0 (PerkinElmer, MA, USA). Lipopolysaccharide (Sigma-Aldrich) was suspended in ultra-pure water.

### Western blot

Western blots were performed as previously described (41). Protein was isolated using M-PER Mammalian Protein Extraction Reagent (Thermo Fisher Scientific), and Halt Protease and Phosphatase Inhibitor Cocktail (Thermo Fisher Scientific). Protein was quantified using the Pierce BCA Protein Assay Kit (Thermo Fisher Scientific), and 10 μg electrophoresed in 4–12% Bis-Tris denaturing gels (Thermo Fisher Scientific) using MOPS buffer chemistry (Thermo Fisher Scientific). Transfer was to 0.2 µm pore PVDF membranes (Bio-Rad Laboratories, CA, USA). Membranes were blocked in 5% skim milk for mouse anti-Actin-β (1:10,000, Sigma-Aldrich, #A1978), rabbit anti-ATG5 (1:2000, AbCam, Cambridge, UK, #ab108327), rabbit anti-Poly-(ADP-ribose) polymerase (**PARP**; 1:3,000, Cell Signaling Technology, Danvers, MS, USA, #9542), mouse anti-NRB1 (1:500, Thermo Fisher Scientific, # H00004077), and rabbit anti-Optineurin (**OPTN**; 1:2000, Cell Signaling Technology, #58981). Blocking with 5% bovine serum albumin (Merck, 126575) was used for rabbit anti-LC3B-I/II (1:3000, Cell Signaling Technology, #4108), rabbit anti-p62/Seqestosome-1 (**p62/SQSTM1**; 1:3000, Cell Signaling Technology, #5114), rabbit anti-TAX1BP1 (**TAX**; 1:2000, Cell Signaling Technology, #5105), rabbit anti-NDP52 (1:4000, Cell Signaling Technology, #60732), rabbit anti-pMTOR^2448^ (1:1000, Invitrogen, MA, USA, # 441125G), and rabbit anti-SIRT-1 (1:1000, AbCam, #ab7343). Membranes were subsequently probed overnight at 4°C with the same respective buffer/antibodies combinations. Secondary incubation was 1 h at ambient temperature in 5% skim milk with mouse or rabbit IgG horseradish peroxidase-conjugated antibodies (R&D Systems, MN, USA, #HAF007 and #HAF008, respectively). The suspension for blocking and antibody incubations was 1× Tris buffered saline (Thermo Fisher Scientific), with 0.1% Tween 20 (Merck). Where possible, outcomes were derived from a single transfer membrane for the respective replicate experiments to minimise inconsistencies produced by inter-blot variation. Chemiluminescent signal production was with Amersham ECL Advanced Western Blotting Detection reagents (GE Healthcare, IL, USA). Image acquisition was performed using the LAS-4000 Luminescent Image Analyzer (Fujifilm Life Sciences, Japan), with signal detection set to annotate blots beyond the linear range (depicted by pink-purple colour). Histogram densitometry was performed using Multi Gauge software (Fujifilm Life Sciences, V3.1, Japan).

### Quantification of IL-6 secretion

IL-6 secretion from THP-1-derived macrophages was quantified in the growth media using enzyme-linked immunosorbent assay (ELISA; Invitrogen, #BMS213INST) following the manufacturer’s protocol.

### Quantification of minimum inhibitory concentration (MIC)

Assessment of MIC was conducted as previously described (37), and in accordance with the Clinical and Laboratory Standards Institute guidelines (42). Briefly, log phase (OD 0.3-0.5, 600 nm) methicillin sensitive *S. aureus* (**MSSA**; ATCC 25923) was diluted in Mueller Hinton broth to a final inoculum of 5 × 10^5^ CFU/well and added to 96-well plates containing a two-fold concentration gradient of antibiotics (0-128 µg/mL). Cultures were incubated for 16 h at 37°C and cell growth quantified (OD 630 nm; t_16_-t_0_ h background subtraction), in a 96-well plate reader (BioTek, Potton Bedfordshire, UK).

### Transmission electron microscopy

PBMC isolates were pelleted (400 g) for 5 min and resuspended/fixed in 500 µL of transmission electron microscopy (TEM) fixative (4% formaldehyde, 1% glutaraldehyde, 4% sucrose in 1× PBS; each Sigma-Aldrich). Fixed cells were processed for mounting by the staff at the University of Adelaide Microscopy suite. Sections were examined using an FEI Tecnai G2 Spirit transmission electron microscope (Thermo Fisher Scientific). For quantitative analysis, autophagic vacuoles were identified using a predefined morphology-based scoring criterion, defined as vesicular structures containing heterogeneous electron-dense cytoplasmic or membranous material and bounded by a limiting membrane. Vesicles that were completely electron-lucent or displayed uniform electron density consistent with lysosomes were excluded from quantification. Vesicle quantification was performed in a blinded manner across treatment conditions.

### Statistical Methods

Statistical analyses were performed using IBM SPSS (v28.0.1.0). A Bayesian gamma (log link) generalised linear mixed model (**GLMM**) was applied (for the respective experiments/outcomes) to account for individual random effects for each sample, and target specific variances. Western blot densitometry results were analysed with a random effect of test repetition and reported relative to the abundance of Actin-β. For the bacterial growth assay, effects of AZM and AZM-[O] on MIC, at each antibiotic concentration, were evaluated. To assess the anti-inflammatory potential of AZM-[O], the effect of treatment*exposure-time interaction on IL-6 secretion was assessed using GLMMs with measurement timepoint (hours) included as a repeated measure. Separate analyses were performed for the 1 and 10 ng/mL LPS-stimulated THP-1 macrophage models.

## Results

### Azithromycin inhibits PBMC autophagy in human blood samples

We first wanted to determine whether AZM is able to modulate autophagy in a model that approximates its administration in humans. To achieve this, we combined pharmacological grade injectable AZM with an *ex vivo* model that enables the organotypic assessment of autophagy in PBMCs derived from human whole blood (38). PBMCs isolated from whole blood exposed to AZM exhibited a significant accumulation of autophagic vacuoles compared to the no-treatment (NT) condition (mean difference +5.566 vs NT, *P* < 0.001), and that was comparable to the influence of CQ (mean difference +4.056 vs NT, *P* < 0.001; **Fig. 1A-E**). While AZM produced a greater effect size than CQ, the difference was not statistically significance (mean difference +1.510, *P =* 0.114). A portion of the PBMCs was also used to assess LC3B-II and p62/SQSTM1 protein abundance. Although the outcomes using protein biochemistry exhibited greater variability per individual (vs quantifying autophagic vacuoles), overall, cells exposed to AZM exhibited a concomitant elevation of LC3B-II and p62/SQSTM1, indicative of restricted autophagy (mean difference in LC3B-II +2.953, *P* < 0.005, and +2.033 for p62/SQSTM1, *P* < 0.001; both vs NT; **Fig. 1F-G**). The effect of AZM matched CQ for the accumulation of LC3B-II, while the abundance of p62/SQSTM1 was marginally higher in PBMCs exposed to AZM (mean difference +0.795, *P* < 0.05, vs CQ). Of note, extending AZM exposure to 4 h – an interval that remains within the organotypic window for autophagy assessment, but beyond the 1 h period originally defined as sufficient for CQ in this system (38), provided an enhanced dynamic range for the protein-based readouts (**Supp. Fig. 1**). Taken together, these findings demonstrate that AZM-mediated restriction of autophagic flux can be detected in human blood samples by protein biochemistry, with TEM providing greater sensitivity and resolution.

**Figure 1:**
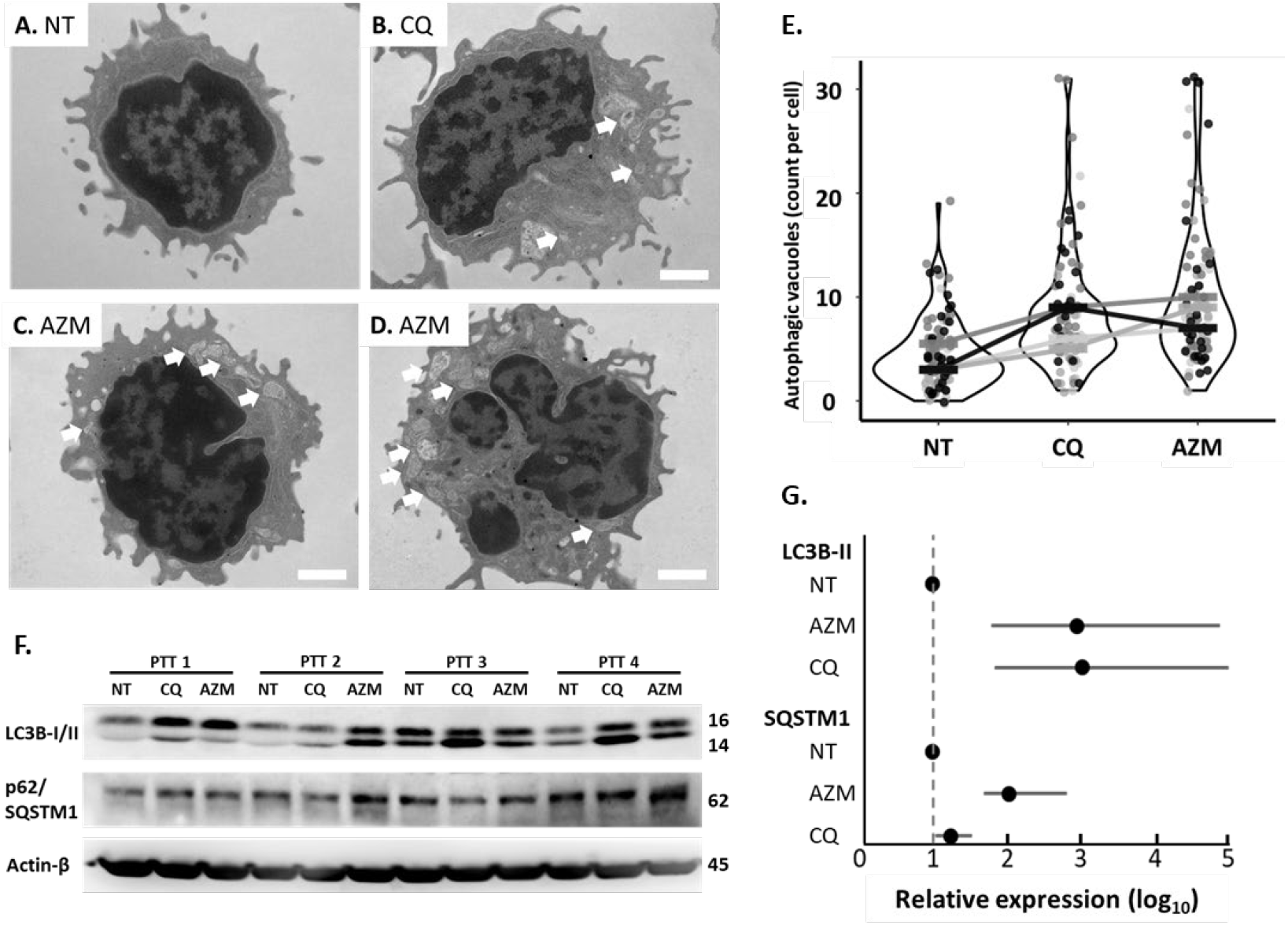
Azithromycin inhibits autophagy in an organotypic intravenous administration model. (**A–D**) Representative transmission electron micrographs of peripheral blood mononuclear cells (PBMCs) under no-treatment (NT) conditions and following exposure to chloroquine (CQ; 150 µM) or azithromycin (AZM; 50 µg/mL) for 1 h, showing accumulation of autophagic vacuoles in treated conditions (as highlighted by arrows). Both AZM (n = 97 cells) and CQ (n = 96 cells) increased autophagic vacuole abundance in PBMCs compared to the NT control (n = 110 cells; *P* < 0.01 for each vs NT). Cell counts were pooled from PBMCs obtained from n = 4 participants. Individual cells are shown as jittered points to reduce overlap. Solid horizontal bars indicate participant-matched median values, with connecting lines denoting within-participant comparisons across treatment conditions. (**F-G**) Parallel protein biochemistry using the same PBMCs samples support a restriction of autophagic flux as evidenced by the concomitant elevation of LC3B-II and p62/SQSTM1 due to AZM that matches the effect of CQ. Outcomes were for n = 4 participants and derived from a single transfer membrane to minimise inconsistencies produced by inter-blot variation. Expression was normalized to Actin-β. Effect sizes represented fold-change relative to the control treatment and uncertainty intervals are ± 95% CI. Results are significant to at least *P* < 0.05 when 95% CI do not cross the dashed line (1-fold).

### The structural modifications required to synthesize azithromycin from erythromycin inhibit autophagy

We tracked the structure-activity relationship of AZM *vs* autophagy in 16HBE14o-cells, in comparison to ERM and two intermediate macrolide derivatives, ERM-oxime and ERM-ether, that are commonly encountered along the synthetic pathway from ERM to AZM (refer to **Fig. 2A** for macrolide structures). Using the abundance of LC3B-II and p62/SQSTM1 as a readout of autophagic flux, we observed a concomitant and progressive increase in the LC3B-II/p62SQSTM1 signal for ERM and ERM-oxime, and a progressive and more significant increase for ERM-ether and AZM. AZM demonstrated the largest effect size for autophagic blockade (i.e. accumulation of LC3B-II +30.7-fold, *P<* 0.001 and p62/SQSTM1 +6.8-fold, *P* < 0.001, *vs* control; **Fig. 2B-C**). Examination of other clinically applied semisynthetic macrolides produced from ERM, including CLM, ROX and TLM, elicited a similar, albeit less pronounced accumulation of LC3B-II and p62/SQSTM1 (*vs* AZM, **Fig. 2B-C**). Observations during routine microscopy showed that each culture/exposure was morphologically comparable to the control, suggesting toxicity was not an obvious contributor to the outcomes for autophagy. For AZM, this was supported by non-significant outcomes for the abundance of cleaved PARP (a product of caspase-3 activity), while the abundance of cleaved PARP due to the other (clinical) macrolides were marginally higher than the observed level of background programmed cell turnover. We also assessed the fourth-generation pre-clinical macrolide SLM, which displayed unique behaviour compared to the other macrolides, causing significant cell death and detachment (cleaved PARP +2.52-fold, *P* = 0.003). The disproportionately high pro-apoptotic effect elicited by SLM is not fully comported here due to signal-loss of cleaved PARP as a result of cell detachment. This is evidenced by a large reduction in its corresponding Actin-β signal (3.23-fold lower than the average Actin-β signal of the other exposures; *P* < 0.001). Only Rubicon and SIRT-1, which have roles in the positive regulation of programmed cell death, produced detectable/consistent signals for the SLM exposure. Notably, and excluding outcomes for SLM, while the expression of LC3B-II and p62/SQSTM1 (effectors of autophagy) demonstrated significant signal modulation (*vs* the control exposure), expression of the upstream pro-autophagy factors p-mTOR^S2448^, Rubicon, SIRT1 and ATG5, were not significantly influenced by either macrolide in the context of this model. Also of note, probing p-mTOR^S2448^ in 16HBE14o-lysates provided two distinct signals of equivalent intensity, at 250 and 288 kDa. Hence, we designated both “p-mTOR^S2448^” because, 1) both are confirmed by the manufacturer(albeit using different cell lysates), 2) related studies (to our knowledge) only report one or the other signal, 3) and we did not observe signal modulation for p-mTOR^S2448^ using the exposures applied here. The significant effect on autophagic blockade observed with AZM, when compared to the other macrolides tested, was attributed to its unique physicochemical characteristics. Specifically, the dibasic nature and net charge of AZM at lysosomal pH values (**Supp. Table 1**) resulting in high lysosomal accumulation through ion trapping (30) is known to induce phospholipidosis (43), and is a likely contributor to autophagic inhibition.

**Figure 2:**
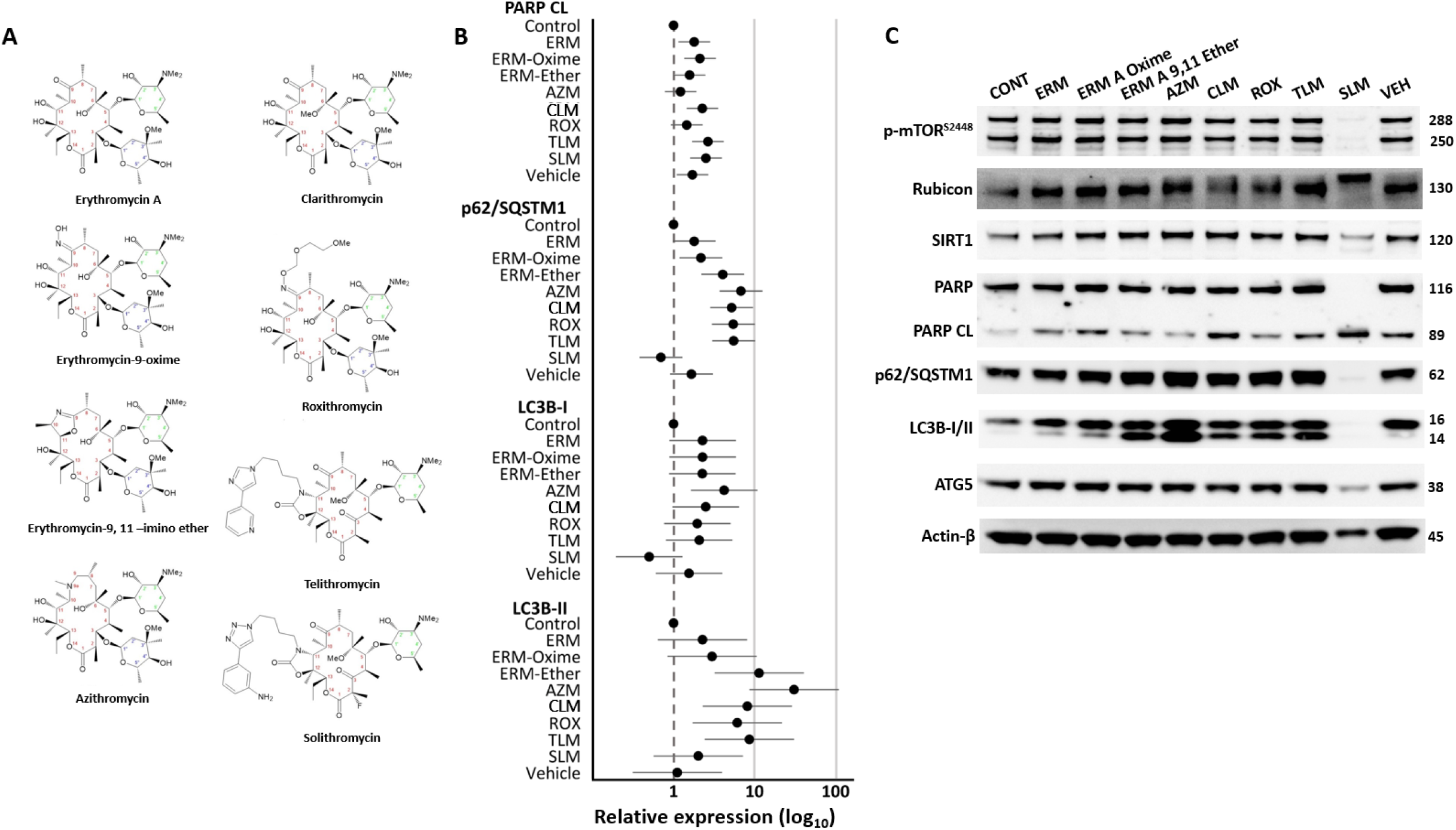
Azithromycin causes a heightened autophagic arrest response in human airway epithelial cells compared to its precursor compounds and related macrolide antibiotics. **A**. Chemical structures of the macrolides. Chemical structures of erythromycin A (**ERM**), erythromycin-9-oxime (**ERM-oxime**), erythromycin-9,11-imino ether (**ERM-ether)**, azithromycin (**AZM**), clarithromycin (**CLM**), roxithromycin (**ROX**), telithromycin (**TEL**) and solithromycin (**SLM)** with systematic numbering of the aglycone lactone, desosamine and cladinose sugars provided in red, green and blue, respectively. The lactone macrocycle (numbering system indicates carbon/nitrogen atoms) is bonded with a desosamine sugar group (upper six-membered ring system attached through C5) for each antibiotic and the cladinose sugar (lower six-membered ring system attached through C3) for each except TEL and SOL. **B**. 16HBE14o-cell responses to the macrolide exposures (16 h, 50 µg/mL). Outcomes represent n = 3 independent cultures/Western blots. Outcomes for the target proteins that demonstrated significant response/modulation to the exposure(s) are shown. Outcomes for each of the respective n = 3 repeated measures of the targets were derived from a single transfer membrane to minimise inconsistencies produced by inter-blot variation. Expression was normalized to Actin-β. Effect sizes represented fold-change relative to the control treatment and uncertainty intervals are ± 95% CI. Results are significant to at least *P* < 0.05 when 95% CI do not cross the dashed line (1-fold). Vehicle (VEH) is ≤ 0.1% DMSO *vs* culture volume. **C**. Western blot showing protein signals derived from the 16HBE14o-model exposed to the various macrolides (16 h, 50 µg/mL). Large and concomitant increase in LC3B-II and p62/SQSTM1 abundance for the AZM exposure (*vs* control), indicates a heightened arrest of autophagic flux (i.e. reduced autolysosomal turnover). This effect appears to be independent of alteration in protein abundance of influential up-stream pro-autophagy factors such as p-mTOR^S2448^ and SIRT1.

### Azithromycin causes the accumulation of autophagy receptors NBR1 and TAX to a greater degree than Optineurin and NDP52

The various autophagy receptor proteins exhibit differential regulation and target specificity (e.g., depolarised mitochondria or intracellular microbes). Hence, we examined Neighbor of BRCA1 gene (**NBR1**), TAX1BP1 (**TAX**), Optineurin, and Nuclear dot protein 52 (**NDP52**), to identify any differential responses to the macrolides, and *vs* observations for p62/SQSTM1. For the treatments in the chemical synthetic pathway from ERM to AZM, the effect size for NRB1 and TAX abundance were strongly influenced by AZM (NBR1 +4.66-fold, *P* < 0.001; TAX +8.68-fold, *P* < 0.001; AZM *vs* control signal; **Fig. 3A-B**). Interestingly for NBR1 (and to a lesser extent TAX; but distinct to the progressive elevation pattern seen in **Fig. 2B** for p62/SQSTM1), the elevation in this particular autophagy receptor was confined to the influences imparted by AZM and CLM, while neither of ERM, ERM-oxime, ERM-ether significantly modulated NRB1 (*vs* the control exposure). In comparison, the abundance of Optineurin and NDP52 remained relatively consistent across each condition, including for the other semisynthetic macrolides CLM, ROX and TLM. Indeed, the most significant influence imparted by CLM, ROX and TLM (when compared to all others) was the upregulation of NBR1. Of note, AZM was not always the most influential exposure; CLM had a greater effect for NBR1, and each of CLM, ROX and TLM increased Optineurin beyond observations for AZM (albeit, neither were significantly larger). Given that the previous outcomes for SLM align with programmed cell death (**Fig. 2B-C**), its influence here for the autophagy receptors (i.e. depletion of a stable/detectable signal), suggests NBR1, TAX, Optineurin and NDP52 do not participate in the positive regulation of apoptosis in the context of this model (at least, as this relates to modulation of their protein abundance). Taken together, these outcomes support a salient effect of AZM to also prolong the duration (and thereby increase the abundance) of autophagy receptors other than p62/SQSTM1 within the cell (*vs* basal conditions in untreated cells). This effect was at least matched by other clinically relevant macrolides for particular autophagy receptors.

**Figure 3:**
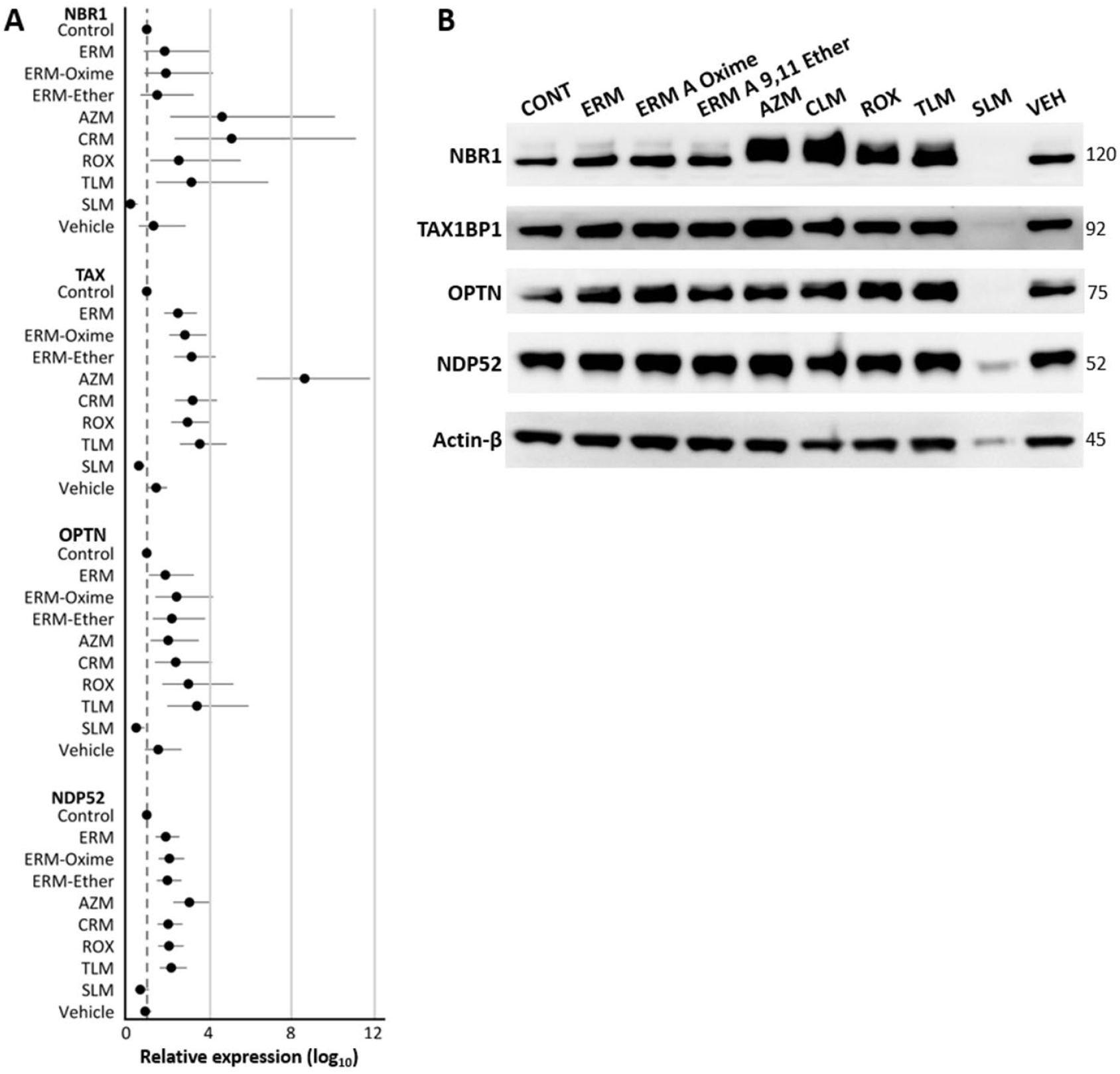
The autophagy receptors NBR1 and TAX are elevated by azithromycin. **A**. Quantification of the protein abundance measures for the autophagy receptors NBR1, TAX, Optineurin and NDP52 in 16HBE14o-cell exposed to the macrolides (16 h, 50 µg/mL). Effect sizes represent n = 3 independent cultures/Western blots. Outcomes for each of the respective repeated measures of the targets were derived from a single transfer membrane to minimise inconsistencies produced by inter-blot variation. Expression was normalised to Actin-β. Effect sizes represented fold-change relative to the control treatment and uncertainty intervals are ±95% CI. Results are significant to at least *P* < 0.05 when 95% CI do not cross the dashed line (1-fold). **B**. Western blot shows autophagy receptor protein signals from 16HBE14o-cultures exposed to the various macrolides (16 h, 50 µg/mL). Responses for NBR1 and TAX show significant signal modulation when the 16HBE14o-cells were exposed to AZM, but relatively less Optineurin (OPTN) and NDP52 was observed to accumulate as a result of this macrolide (*vs* the control exposure). Interestingly, CLM elicited a stronger (but not statistically distinct) effect for the elevation of NBR1 *vs* AZM, and each of CLM, ROX and TLM produced a similar outcome (*vs* AZM) for the autophagy receptor Optineurin.

### Modification of azithromycin’s charged state significantly reduces its arrest of autophagy

One mechanism for AZM inhibition of autophagy is entry into lysosomes where it becomes strongly protonated, depleting the local hydronium concentration, and leading to an increase in pH. Protonation of both the 9a-lactone and 3’-desosamine amine groups resulting in a double net positive charge effectively ‘traps’ AZM enabling this lysosomotropic effect. To confirm this mechanism contributes to the arrest of autophagy, we synthesised AZM-[O] by oxidising the desosamine group, thereby eliminating its ability to become protonated and therefore reducing the overall net charge of the molecule *vs* AZM (**Fig. 4A; Supp Table 1**). At the highest test concentration (50 µg/mL), oxidation of the desosaminyl amine appreciably lowered the accumulation of both LC3B-II and p62/SQSTM1 in 16HBE14o-cells (AZM-[O] *vs* AZM, -22.07-fold and -2.88-fold, respectively, both *P* < 0.001; **Fig. 4B-C**). Examining the dose titration, both compounds elicited similar effects for the accumulation LC3B-II and p62/SQSTM1 for the 2.5 µg/mL intervention but became distinct at five-times this amount (10 µg/mL, *P* > 0.05). Of note, here the observations for cleaved PARP (apoptosis induction) grouped to produce an overall elevated outcome (*vs* outcomes in **Fig. 2**), for 10 and 50 µg/mL of AZM (2.31-fold, *P* = 0.004 and 3.96-fold, *P* < 0.001, respectively, *vs* control). While this agrees with inhibiting the pro-survival influences imparted by autophagy, routine microscopic observations showed the cells were not morphologically distinct from the control exposure.

**Figure 4:**
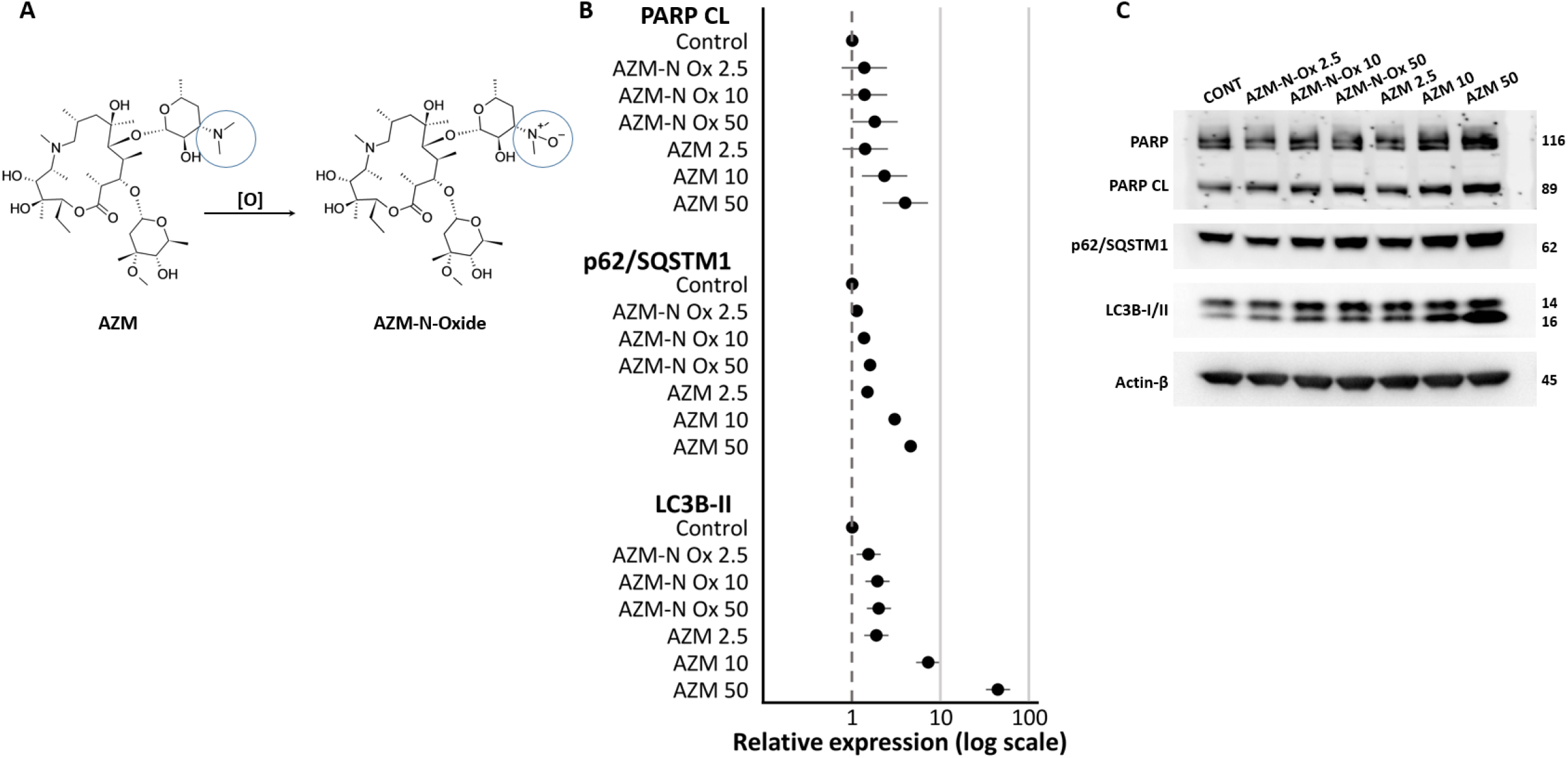
Structure-activity relationship of azithromycin vs autophagy shows oxidation of the desosaminyl amine diminishes its off-target block of autophagy. **A**. Scheme showing oxidation of AZM desosaminyl amine. Oxidation of the desosaminyl amine (blue circles) produces a zwitterionic *N*-oxide group (no net charge), blocking the potential protonation of the amine group. **B**. Quantification of the outcomes of Western blot analyses shows the concomitant increase in the abundance of LC3B-II and p62/SQSTM1 for AZM is significantly reduced for the AZM-[O] derivative. The outcomes for AZM and LC3B-II / p62/SQSTM1 follow a dose relationship (2.5, 10, and 50 µg/mL; 16 h). Here, this outcome is matched by a progressive increase in apoptosis with AZM (as assessed via the abundance of cleaved PARP), which is not observed after exposure with the oxidised form. Expressions was normalized to Actin-β. Effect sizes represented fold-change relative to the control treatment and uncertainty intervals are ±95% CI for n = 3 experiments. Results are significant to at least *P* < 0.05 when 95% CI do not cross the dashed line (1-fold). **C**. Representative Western blot shows the processing of PARP to produce cleaved PARP, and the concomitant progressive elevation of LC3B-II and p62/SQSTM1 in 16HBE14o-cells after stimulation with increasing concentrations of AZM, but which is not observed for AZM-[O]. Actin-β served as the loading control.

### Oxidation of the AZM desosamine group reduces its bacteriostatic activity

We assessed the MIC for dose titrations of AZM and AZM-[O] to determine whether the oxidation of AZM desosaminyl amine modulates the bacteriostatic activity observed for AZM. Shown in **Fig. 5**, AZM exhibited exceptionally effective bacteriostatic activity, inhibiting the growth of the MSSA reference strain at an MIC within 1-4 µg/mL. In contrast the antimicrobial activity of AZM-[O] was significantly attenuated and was only able to affect a bacteriostatic outcome at the maximum exposure of 128 µg/mL. This outcome is consistent with the essential function of the tertiary amine of the AZM desosamine group as an important structural determinant that enables its inhibition of the bacterial 50S ribosomal subunit.

**Figure 5:**
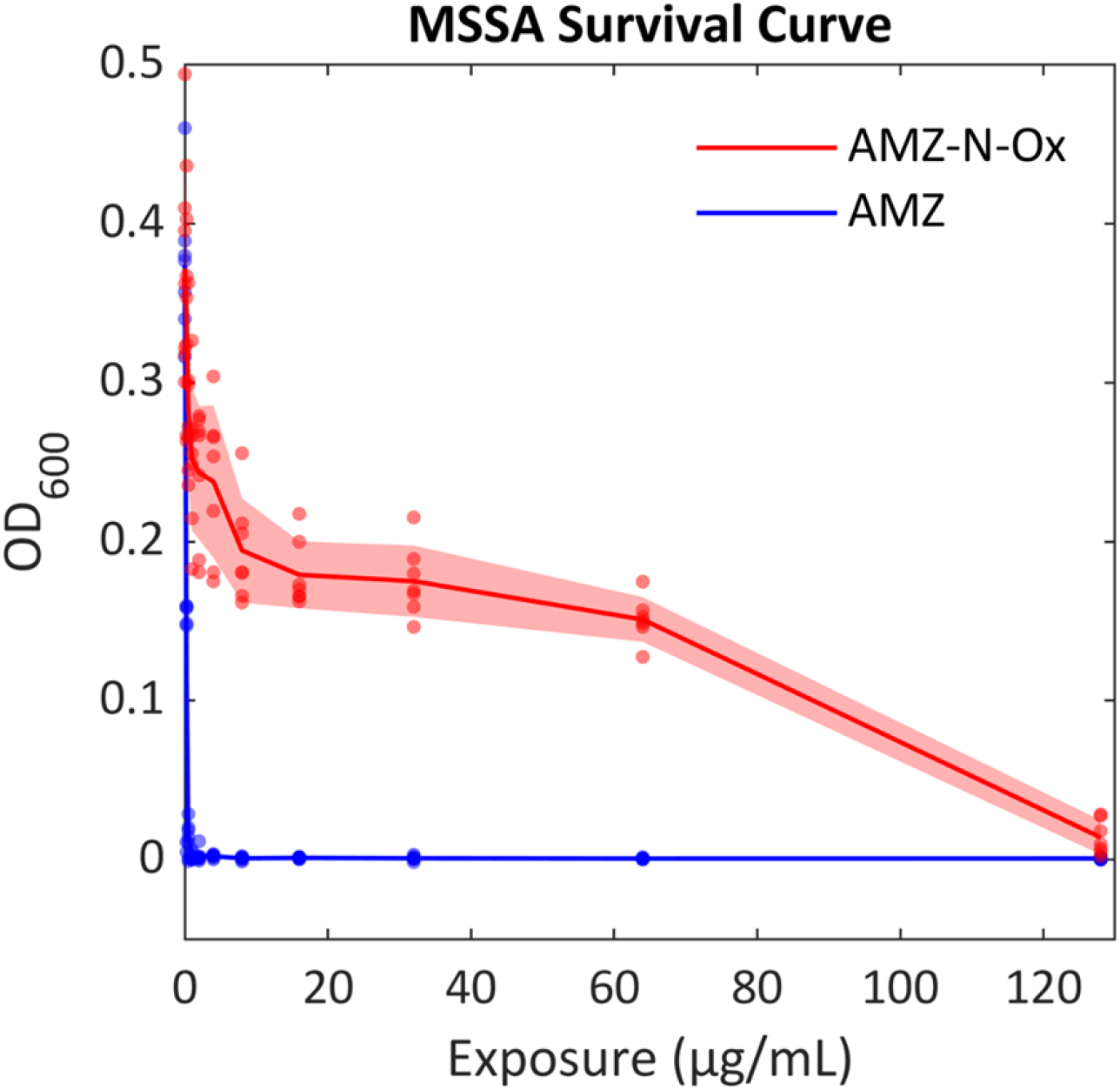
AZM-[O] exhibits low bacteriostatic efficacy vs the AZM parent molecule. MSSA was used to model the comparative bacteriostatic potential of AZM *vs* the AZM-[O] derivative. AZM was extremely effective in preventing the growth of MSSA (MIC of ∼ 2 µg/mL over a 16 h assay period), whereas AZM-[O] approached MIC at the maximum dose of 128 µg/mL (over 64-fold reduction in bacteriostatic activity *vs* AZM). The bacteriostatic effect of AZM-[O] was approximately 105% lower than AZM at every exposure dose (*P* < 0.001). Outcomes represent n = 7 replicate experiments; shaded area represents ±SD.

### AZM-[O] exhibits attenuated anti-inflammatory properties *vs* AZM

One beneficial off-target influence imparted by AZM is a moderate anti-inflammatory effect. To determine whether AZM-[O] retained this anti-inflammatory property, we quantified the magnitude of secreted IL-6 in an LPS-stimulated THP-1 macrophage model, co-exposed to either AZM or AZM-[O]. AZM significantly moderated the secretion of IL-6 in THP-1 macrophages treated with 1 ng/mL LPS for the 6 and 24 h exposure intervals (-95.80 ng/mL, *P* < 0.001 and-86.19 ng/mL, *P* = 0.002, *vs* control, respectively; **Fig. 6A**), while AZM-[O] elicited a similar, albeit less influential effect (-22.55 ng/mL, *P* < 0.04 and -55.37 ng/mL, *P* = 0.04, *vs* control, respectively). In contrast, outcomes for AZM-[O] with 10 ng/mL LPS for either time interval were indiscernible from the control exposure (**Fig. 6B**). Interestingly, AZM was also less effective in the 10 ng/mL LPS model, with a significant outcome for the 6 h LPS exposure (-130.43 ng/mL, *P* < 0.001, *vs* control), but a non-significant result for the 24 h exposure (-62.9 ng/mL, *P* = 0.161, *vs* control). That AZM was also relatively less effective for the 24 h interval with 1 ng/mL LPS (*vs* the 6 h interval), hints to a washout mechanism possibly related to chemical stability within the cell, and/or the turnover of organelles and secretion of vesicles containing AZM.

**Figure 6:**
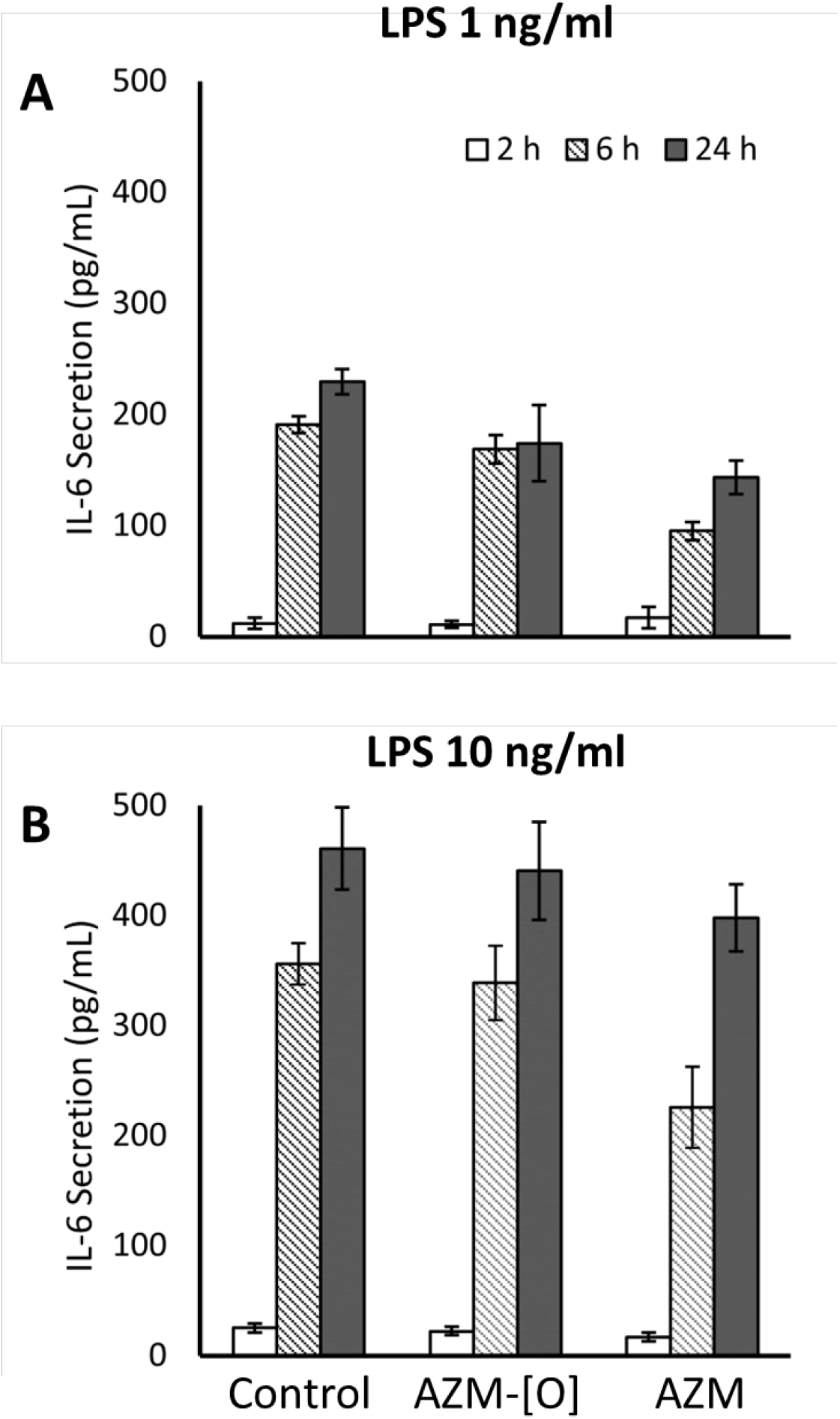
Oxidation of azithromycin’s desosaminyl amine attenuates its anti-inflammatory activity. The secretion of IL-6 was used to assess the anti-inflammatory effect imparted by AZM *vs*. AZM-[O]. THP-1 macrophages stimulated with LPS respond by secreting IL-6 into the media, particularly following 6 and 24 h incubation. AZM’s reduction of IL-6 was significant (vs control) after co-exposure with 1 ng/mL LPS for the 6 and 24 h treatment (44)also elicited a significant, albeit attenuated (vs AZM), reduction of IL-6 for 1 ng/mL LPS after the 6 and 24 h exposure intervals (*P* < 0.04 and *P* = 0.04, respectively), while outcomes for AZM-[O] using 10 ng/mL LPS were comparable to the control exposure. Outcomes represent n = 3 separate experiments; confidence intervals are ±SD.

## Discussion

The preservation of frontline antibiotics against increasingly resilient bacterial pathogens is a global health imperative. While there remains an apparent lack of urgency to develop new antibiotics (35), an equally important challenge is the absence of predictive models capable of identifying clinically relevant off-target effects, as highlighted by the failure of the promising macrolide-ketolide SLM (45). For AZM, this challenge is compounded by the coexistence of deleterious off-target effects, such as mitochondrial toxicity (46), alongside beneficial immunomodulatory properties that underpin its widespread and prolonged use. We predict that this has obscured gaps in our understanding of its impact on fundamental host-cell processes. Alternatives such as ERM lack the well described immunomodulatory properties that underpin the clinical benefit of long term macrolide therapy (44), and is less well tolerated during prolonged administration (47). In the context of chronic airway disease, such use creates a scenario which inadvertently contribute to both disease persistence and the gradual erosion of AZM’s long-term clinical utility. There are now increasing calls to develop regulatory regimes specific to AZM, to combat the emergence of AZM tolerant species (e.g., (48)). There is even less information warning of AZM’s potent off-target inhibition of autophagy - a fundamental survival process that also serves as the cells’ bacterial clearance mechanism. Chronic respiratory diseases like COPD, cystic fibrosis (CF) and non-CF bronchiectasis are driven by bacterial infections, and each are linked to dysregulation of autophagy (49). Here we demonstrate for the first time that AZM potently inhibits autophagy in human tissues. We also showed that targeting its charged state (AZM-[O]) at the desosaminyl amine is one method to overcome this off-target effect.

While we and others have previously shown that AZM restricts autophagy in cell models, to our knowledge this phenomenon has not been convincingly demonstrated in clinically-derived tissues. Indeed, even *in vitro* models require substantial experimental manipulation and orthogonal measures to enable meaningful assessments of autophagic flux (50). Consistent with this, direct quantification of autophagic flux from human tissue samples has remained largely inaccessible, especially as cells/autophagy rapidly respond to alterations in external conditions (51). To overcome this, we applied a recently developed *ex vivo* approach that enables organotypic assessment of autophagy in PBMCs maintained within whole blood (38). This system preserves basal autophagic function outside the body, permits controlled pharmacological exposure, and closely approximates the clinical scenario of systemic AZM administration. Using this model, TEM showed an accumulation of autophagic vacuoles in AZM-treated PBMCs, consistent with restricted autophagosome turnover. This outcome closely phenocopied observations with CQ, which has similar lysosomotropic properties and its capacity for ion trapping within the lysosome, leading to impaired autophagosome– lysosome fusion (32). These ultrastructural findings were corroborated by protein biochemical analyses demonstrating concomitant accumulation of LC3B-II and p62/SQSTM1. As expected, effect sizes derived from Western blotting were less pronounced, reflecting the semi-quantitative nature of this approach and reliance on secondary signals from individual factors involved in autophagy (vs directly quantifying autophagosomes). However, extending AZM exposure beyond the protocol’s 1 h interval (which was optimal for CQ (38, 52)) to 4 h, increased the dynamic range of the readouts relative to baseline (**Supp. Fig. 1**). Collectively, these findings demonstrate that AZM is an influential inhibitor of autophagic flux in a human PBMCs, and given the organotypic design of this model, we predict this phenomenon also occurs in patients who are administered AZM.

Comparison of clinically used macrolides with intermediate compounds involved in AZM synthesis, revealed SAR that influence autophagic flux. While the accumulation of AZM in the lysosome resulting from ion trapping of its doubly charged (2+) state correlates with autophagic inhibition, other macrolides also display upregulated levels of markers of autophagic arrest despite very different protonation states. Excluding SLM for obvious toxicological reasons, ERM-ether and TLM are the only other macrolides tested that have ionisable basic groups other than the desosamine group, which is common to all of the macrolides. Thus, the decrease in lysosomal pH relative to the cytosol could lead to an increase in the net charge (> 1+) of ERM-ether and TLM resulting in their retention (**Supp. Table 1**), which may contribute to an increase in lysosomal pH and autophagic arrest, as observed for AZM. This is somewhat supported by the similarly high accumulation of TLM and AZM previously reported in polymorphonuclear neutrophils (53-55). In comparison, the accumulation of CLM and ROX in polymorphonuclear neutrophils is significantly lower, which is consistent with their lower net charge (≤ 1+). Nevertheless, CLM and ROX displayed upregulation of LC3B-II and p62/SQSTM1 similarly to TLM, suggesting an alternative (and/or an additional) mechanism of autophagic arrest for these macrolides. Unfortunately, calculation and comparison of the physiochemical properties of the macrolides did not provide further insights that would explain these observations (**Supp. Table 1**).

Related to this, we noted that the autophagy receptors exhibited differential abundance profiles, depending on which macrolide was applied to the model. This outcome, coupled with the protracted duration (days) that macrolides can remain within the cellular environment, means they have an extended opportunity to perturb further and distinct autophagy-related factors beyond the arrest of autophagosome-lysosome fusion (54, 55). One possibility is that the macrolide-dependent differences arise due to distinct influences on the factors that govern the unique post-translational modifications observed for the various autophagy receptors (56). More generally, they may modulate the numerous up-stream regulators of the autophagy pathway (e.g. 51), and/or the overlapping cellular processes that influence autophagy during (for example) innate immunity and cells survival (noted here with our observations for PARP/apoptosis). To begin addressing these mechanistic complexities, we developed a photoaffinity labelled AZM probe capable of covalent crosslinking to proximal cellular components. Preliminary data demonstrate dose dependent and competitively displaceable engagement of cellular proteins in airway epithelial cells, consistent with specific AZM interactions (**Supp. Fig. 2**). While identification of individual targets is beyond the scope of the present study, these findings support a protein mediated mechanism underlying AZM’s influence on the host cell (57, 58). Ongoing work aims to define the AZM protein interactome, inform structure activity relationships for new AZM derivatives, and functionally validate compounds that mitigate autophagy arrest while preserving the immunomodulatory properties of the parent molecule.

One such AZM derivative we assessed here was AZM-[O]. We predicted AZM-[O]’s altered charge state (vs AZM) would mitigate the off-target inhibition of autophagy observed for the parent molecule. While this outcome was realised, this particular modification (oxidation of the desosaminyl amine) also significantly diminished the antibacterial and anti-inflammatory activities observed with AZM. An ideal outcome would have been a derivative that corrected the off-target block of autophagy, was non-antibiotic, but that retained its immunomodulatory activity. Such a derivative would provide a novel non-steroidal anti-inflammatory that concentrates at the airways, without contributing to the selection of AZM-resistance bacteria (59). The challenge here is that AZM’s unique anti-inflammatory effect(s) await a clear description (60), and we suspect these properties are mediated (and hidden amongst) intricate cell-cell interactions at the organ level, e.g., during the type of inflammation synonymous with COPD and CF. For autophagy, AZM has been noted to indirectly influence the mTOR pathway (via inhibition of S6RP phosphorylation) in the context of idiopathic bronchiolitis, albeit with a high concentration of AZM (300 µg/mL) (61). However, no studies have attempted to identify specific lysosomal ion channel or transporter binding domains that could inform SAR studies to address AZM-mediated suppression of autophagosome-lysosome fusion. The most promising nonantibiotic AZM derivative reported thus far (to our knowledge) was by Kragol et al. (2022), who synthesized a stereoisomer of AZM that retained significant immunomodulatory activity, albeit the suppressive effects on autophagy were not examined (62). Nevertheless, the combination of this outcome with ours for AZM-[O] provide promising evidence that the structural determinants of AZM’s antibacterial activity, and the off-target effects for autophagy and innate immunity, may be mutually exclusive.

In summary, we showed AZM reduces autophagy flux *ex vivo* and defined SAR that underpins AZM-mediated inhibition of autophagic flux. These findings highlight that AZM exert previously underappreciated effects on fundamental host-cell processes. We predict these effects drive the decline in the clinical utility of AZM therapy observed during the management of chronic respiratory disease. Systematic identification of such off-target activities is therefore essential, both to inform safer use of existing antibiotics and to guide the rational development of next-generation compounds with improved host compatibility.

## Acknowledgements

We are grateful for the expertise and support of the Royal Adelaide Hospital’s SA Pathology phlebotomy team and Thoracic Department, and the participants enrolled into this study who generously provided their valuable time and tissue samples. We are also thankful for: cytological support from Prof Angel Lopez (SA Pathology); pharmacological insight provided by Prof Benedetta Sallustio (Adelaide University) and Dr Sally Marotti (Adelaide University and SA Health); assistance in chemical synthesis by Christian Danker (Adelaide University); Sean Quang Doan who assisted with the transmission electron microscopy. This study was funded by the Royal Adelaide Research Committee, Royal Adelaide Hospital Research Fund, The Health Services Charitable Gifts Board (both the SA Pathology Division and Royal Adelaide Hospital Division), and the Rebecca L Cooper Medical Research Foundation.

## Supplementary Figure Legends

***supp Fig. 1. LC3B-II protein signals were progressively elevated in whole blood-derived PBMCs in response to increasing periods of AZM exposure***.

(**A-B**) Untreated (“no treatment”, NT) blood samples spanning the 4 h exposure interval exhibited similar base-line levels of LC3B-II abundance in PBMCs. However, LC3B-II signals elevated with time (*vs* either NT control) in the presence of AZM (*P* < 0.05 minimum, for each comparison of AZM exposure vs either NT condition, with the exception of NT 4 h *vs* AZM 1 h; student t-test). Outcomes are from n = 3 separate experiments using one blood donor. Outcomes were derived from a single transfer membrane to minimise inconsistencies produced by inter-blot variation. A common calibrator (“Calibrt’n”) sample was used to account for intra-blot variation across the triplicate series. Expression was normalized to Actin-β.

***Supp Fig. 2. Signals indicative of AZM-protein interactions from 16HBE14o-airway epithelial cells exposed to our custom azithromycin photoaffinity labelled probe***.

*In situ* application of our photoaffinity labelled (PAL) AZM probe, which incorporates a diazirine crosslinker and terminal alkyne handle for downstream bioorthogonal ligation (hereafter referred to as AZM PAL), to the 16HBE14o-model. Cells were exposed to AZM PAL (4 h) followed by UV-irradiation to enable covalent cross-linkage to proteins proximal to AZM PAL. (**A-C**) Exposures/lane: lane 1 – DMSO vehicle (veh); lanes 2-4 – increasing concentrations of AZM PAL; lane 5 – cells were pre-exposed for 1 h to AZM (200 µM) to block potential binding partners prior to exposure with AZM PAL; lane 6 – no click (“-click”) control to prevent fluorescein biotin azide (BroadPharma; #BP-28017) binding to the alkyne handle of AZM PAL. Results show a dose-dependent increase in either FITC (SDS-PAGE) or biotin interaction signals (NeutrAvidin Protein probe; Thermo Fisher Scientific, #31000; Western blot; A-B). Pre-incubation with AZM reduced either signal elicited by the 10 µM AZM PAL exposure (i.e. lane 3 vs 5). Total protein loading (and molecular weight markers) was assessed using Coomassie blue staining of the SDS-PAGE gel (C).

