## Supplemental Figs for "Modifying Azithromycin to Mitigate Arrested Autophagy"

Supp Fig 1

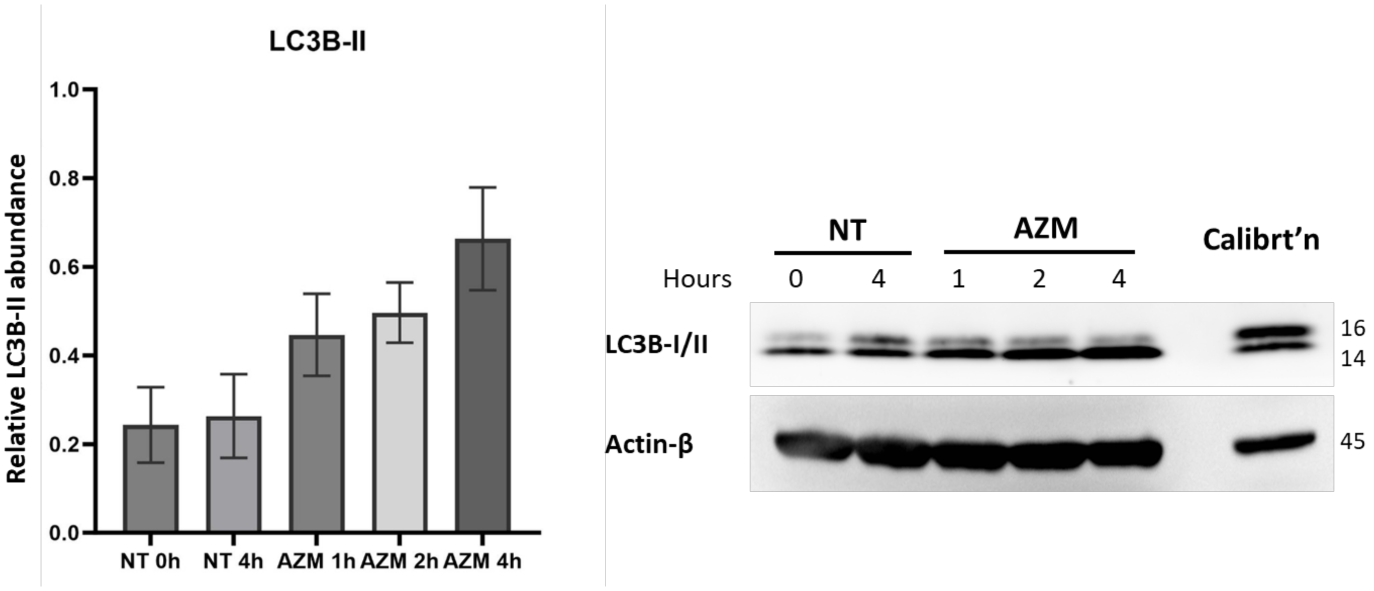
LC3B-II
Relative LC3B-II abundance
1.0
0.8
0.6
0.4
0.2
0.0
NT 0h
NT 4h
AZM 1h
AZM 2h
AZM 4h
NT
AZM
Calibrt'n
Hours
0
4
1
2
4
LC3B-I/II
Actin-β
16
14
45

Supp Fig 2

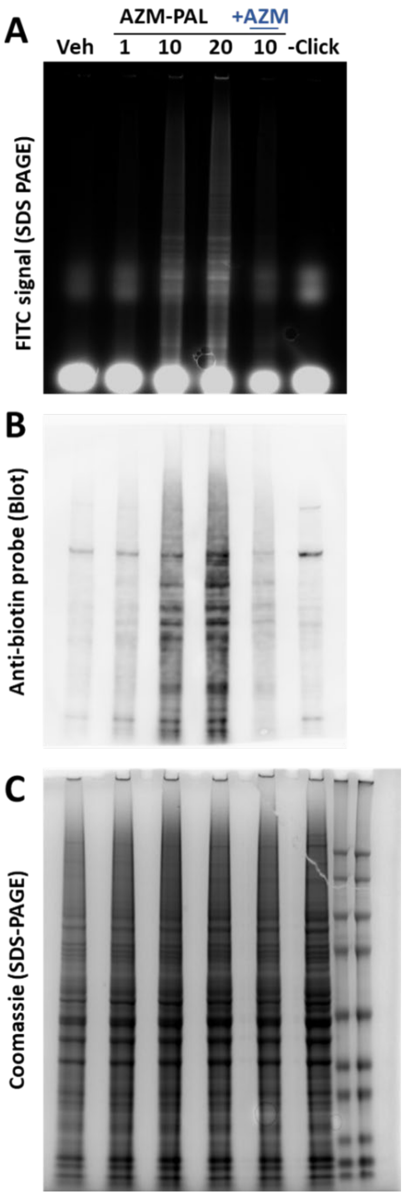
A
AZM-PAL
+AZM
Veh
1
10
20
10
-Click
FITC signal (SDS PAGE)
B
Anti-biotin probe (Blot)
C
Coomassie (SDS-PAGE)

### Supp Table 1

| Macrolide | $pK_a$ values[a] | Net Charge[b] | | LogP[c] | Log D[c] | | TPSA[c] | Molar reflectivity[c] | Polarisability[c] | #HBA[c] | #HBD[c] | I/E ratio[d] |
|---|---|---|---|---|---|---|---|---|---|---|---|---|
| | | pH 7.4 | pH 5.0 | | pH 7.4 | pH 5.5 | (Å$^2$) | (m$^3$.mol$^{-1}$) | (Å$^3$) | | | |
| ERM | 8.88 | 0.97 | 1.00 | 2.83 | 1.69 | -0.02 | 194 | 189.2 ± 0.4 | 75.0 ± 0.5 | 14 | 5 | 13-25 |
| ERM-oxime | 8.88 | 0.97 | 1.00 | 3.04 | 2.11 | 0.40 | 209 | 185.9 ± 0.5 | 73.7 ± 0.5 | 15 | 6 | |
| ERM-ether | 8.88, 5.5 | 0.98 | 1.76 | 2.67 | 2.17 | -0.57 | 178 | 183.1 ± 0.5 | 72.6 ± 0.5 | 14 | 4 | |
| AZM | 9.45, 8.74 | 1.95 | 2.00 | 3.33 | 1.36 | -0.79 | 180 | 197.6 ± 0.4 | 78.3 ± 0.5 | 14 | 5 | 250-387 |
| CLM | 8.99 | 0.97 | 1.00 | 3.16 | 2.38 | 0.67 | 183 | 194.0 ± 0.4 | 76.9 ± 0.5 | 14 | 4 | 100 |
| ROX | 9.17 | 0.98 | 1.00 | 3.73 | 2.80 | 1.10 | 217 | 207.5 ± 0.5 | 82.3 ± 0.5 | 17 | 5 | 90 |
| TLM | 8.7, 5.1, 3.1 | 0.96 | 1.57 | 4.52 | 3.62 | 1.87 | 172 | 216.2 ± 0.5 | 85.7 ± 0.5 | 15 | 1 | 275 |
| SLM | 9.44, 3.94 | 0.99 | 1.08 | 3.44 | 2.90 | 1.19 | 198 | 218.5 ± 0.5 | 86.6 ± 0.5 | 16 | 3 | |
| AZM-[O] | 9.45 | 0.99 | 1.00 | | | | | | | | | |

[a] ERM, AZM and CLM $pK_a$ values determined by potentiometric titration, J. McFarland *et al.*, *J. Med. Chem.* **1997**, 49(9), 1340; ROX $pK_a$ value determined by potentiometric titration, Z. Qiang *et al.*, *Water Res.* **2004**, 38, 2874; TLM $pK_a$ values reported by D. Vazifeh *et al.*, *Antimicrob. Agents Chemother.* **1998**, 42(8), 1944; SLM $pK_a$ values reported by D. Evans *et al.*, *J. Pharm. Sci.* **2017**, 107(1), 412; ERM-oxime $pK_a$ value presumed to be similar to ERM; ERM-ether desosamine $pK_a$ value presumed to be similar to ERM, while iminoether motif estimated to have a $pK_a$ value (5.5) resembling structurally analogous oxazolines, G. R. Porter *et al.*, *Nature* **1958**, 4640, 927; AZM-[O] $pK_a$ value presumed to be similar to AZM 9a-amine. [b] Net charge calculated from $pK_a$ values. [c] Physiochemical properties determined in ChemSpider (Royal Society of Chemistry, ChEBI release 150) with the ACD/Labs Percepta Platform plugin (PhysChem Module, version 14.00); TPSA = topological polar surface area; #HBA = number of hydrogen-bond acceptors; #HBD = number of hydrogen-bond donors. [d] Intracellular-to-extracellular concentration ratio (I/E) of macrolides in human polymorphonuclear neutrophils *in vitro* following 2 h of incubation; values taken from M. T. Labro, *Clin. Microbiol. Infect.* **1996**, S24, W. L. Hand *et al.*, *Int. J. Antimicrob. Agents* **2001**, 18(5), 419 and D. Vazifeh *et al.*, *Antimicrob. Agents Chemother.* **1998**, 42(8), 1944.
